# Simultaneous Functional Ultrasound, Intrinsic Optical Signal and Widefield Calcium Neuroimaging

**DOI:** 10.64898/2026.06.27.733865

**Authors:** Shubham Mirg, Prameth Gaddale, Achutha Kumar, Krishnendu Samanta, Bhawna Saini, Saniya P. Patil, Ana A. Vargas, Aayushi Laliwala, Agata A. Exner, Yong Wang, Grayson O. Sipe, Sri-Rajasekhar Kothapalli

## Abstract

Functional ultrasound (fUS) maps cerebral blood volume (CBV) but lacks molecular and neuronal specificity. By simultaneously integrating fUS with optical imaging, we show that fUS-derived CBV correlates with both optically measured hemoglobin and neuronal calcium activity in awake mice. We further derive hemodynamic response functions linking calcium activity to CBV during spontaneous and sensory-evoked activity. Application to a mouse glioblastoma model demonstrates utility for studying neurovascular dysfunction in complex neuropathologies.

## Main

Functional ultrasound (fUS) imaging maps brain-wide cerebral blood volume (CBV) changes at high spatial (~100 μm) and temporal resolutions (1–10 Hz)^1^. fUS has been rapidly adopted across preclinical and clinical applications, including rodent and non-human primate studies, neonatal and adult human imaging, and brain-computer-interface (BCI) applications^2–5^. The fUS-derived CBV signal originates from Doppler frequency shifts in backscattered echoes produced by moving red blood cells. Yet, like fMRI, fUS remains an indirect surrogate for neuronal activity, motivating widespread interest in characterizing its precise neurovascular origins.

To interrogate this relationship, previous studies have combined fUS with electrophysiology^6^. Notably, simultaneous Neuropixels recordings and fUS revealed that arousal-dependent, opposing excitatory and inhibitory neural populations differentially drive fUS-derived CBV changes^7^. While such recordings provide high temporal resolution and deep-brain access, they lack cell-type and molecular specificity. Optical and photoacoustic techniques offer a non-invasive alternative with direct molecular access. However, prior studies integrating optical imaging with fUS remain limited to sequential measures^8,9^, precluding direct temporal correspondence and restricting analyses to stimulus-evoked conditions. Our prior interleaved photoacoustic-fUS work showed agreement between oxygen saturation and CBV changes during hypercapnia, but lacked the high frame-rate (~0.3 Hz) and true simultaneity^10^. Concurrent fUS-optical imaging can overcome each modality’s limitations: optical imaging resolves cortex-wide molecular dynamics but only superficially, while fUS offers whole-brain depth without molecular specificity. Combining them answers fundamental questions neither modality can address alone: how deep hemodynamics relate to cortical neuronal activity, what molecular processes underlie the fUS signal, and how neuronal calcium maps onto hemodynamic changes through cross-modal transfer functions.

Nevertheless, to our knowledge, simultaneous acquisition of fUS, optical hemoglobin dynamics, and widefield neuronal calcium activity has not been demonstrated. Technical challenges persist due to the optical opacity of conventional ultrasound transducers. While photoacoustic imaging utilizes acousto–optic combiners^11^ to permit simultaneous optical illumination and acoustic detection, comparable strategies have not been demonstrated for combining fUS with optical neuroimaging. Here, using a custom, compact acousto-optic combiner, we integrated intrinsic optical signal imaging (IOSI) and widefield calcium fluorescence imaging to simultaneously measure hemoglobin and neuronal dynamics alongside fUS-derived CBV changes, in both awake and anesthetized mice.

Our fUS-optical imaging approach is presented in Fig. 1a–c (Supp. Fig. 1; Methods). The system resolved optical features to ~5 μm (Supp. Fig. 2), suitable for microscale, cortex-wide optical imaging. We first performed simultaneous fUS-optical imaging through a chronic cranial window in awake, head-restrained transgenic mice expressing GCaMP8s under the CaMKIIα promoter (Supp. Video 1). We obtained baseline-normalized CBV changes (ΔCBV-fUS) from fUS imaging; baseline-normalized oxy-, deoxy-, and total hemoglobin (ΔHbO, ΔHbR, ΔHbT) from interleaved 530 and 625 nm reflectance IOSI images; and neuronal calcium (ΔCa^2+^) from 470 nm GCaMP fluorescence images (Fig. 1d–f; Supp. Video 2; Methods).

**Figure 1:**
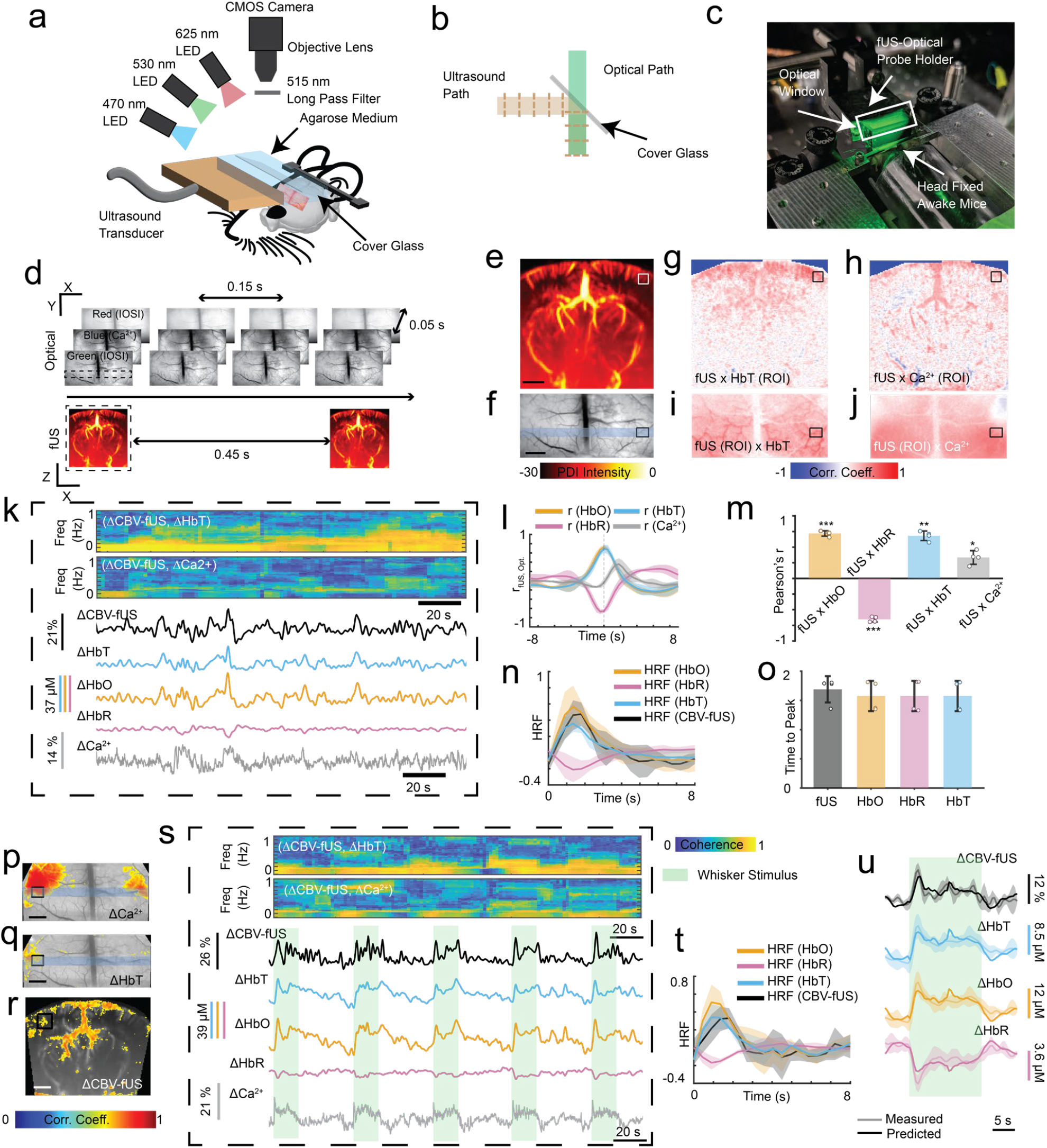
Simultaneous fUS-optical imaging of awake mouse brain activity. **(a)** Combined fUS-optical imaging setup. **(b)** Co-aligned optical and ultrasound paths via a 45°-oriented cover-glass reflector. **(c)** Awake mouse fUS-optical imaging setup. **(d)** Timing diagram for fUS and multi-wavelength optical frames. **(e, f)** Simultaneously acquired fUS-derived cerebral blood volume (CBV) and optical image; blue patch marks the fUS plane. **(g, h)** Pixelwise correlation of baseline-normalized CBV (ΔCBV-fUS) maps with averaged baseline-normalized total hemoglobin (ΔHbT) and calcium (ΔCa^2+^) over the ROI in (f). **(i, j)** Reverse-direction pixelwise correlation of ΔHbT and ΔCa^2+^ (~1.5 s delayed) maps with averaged ΔCBV-fUS over the ROI in (e). **(k)** Coherogram of ΔCBV-fUS with ΔHbT, and ΔCa^2+^ and filtered traces derived from the ROIs in (e, f). **(l)** Cross-correlation of ΔCBV-fUS with ΔHbO, ΔHbR, and ΔHbT and ΔCa^2+^ (n = 4). **(m)** Pearson’s correlation for ΔCBV-fUS with ΔHbO, ΔHbR, and ΔHbT and ΔCa^2+^. **(n)** Hemodynamic response functions (HRFs) linking ΔCa^2+^ to ΔCBV-fUS, ΔHbO, ΔHbR, and ΔHbT (n = 4). **(o)** HRF time-to-peak, indicating neurovascular coupling delay. **(p, q)** Optical and **(r)** fUS images overlaid with whisker-evoked ΔCa^2+^, ΔHbT, and ΔCBV-fUS correlation maps (r > 0.6, p<0.05). **(s)** Coherogram of ΔCBV-fUS with ΔHbT, and ΔCa^2+^ and filtered traces derived from the ROIs in (p-r). **(t)** HRFs from individual whisker trials per signal (n = 3, 5 trials each). **(u)** Predicted traces overlaid with averaged whisker responses. Scale bars, 1 mm. *p < 0.05, **p < 0.01, ***p < 0.001. (PDI: Power Doppler Intensity)

Next, leveraging simultaneous fUS and optical measurements, we assessed cross-modal correspondence in two directions. First, seeding the ROI-averaged ΔHbT signal (bandpass 0.02– 0.5 Hz) and cross-correlating it pixelwise against the ΔCBV-fUS map localized the strongest fUS correlations to the same cortical region as the optical seed (Fig. 1g); seeding the high-pass-filtered ΔCa^2+^ signal (>0.02 Hz, delayed ~1.5 s) yielded the same co-localization (Fig. 1h). Reversing the direction, by seeding ΔCBV-fUS and mapping onto the ΔHbT and ΔCa^2+^ images, reproduced this spatial correspondence (Fig. 1i, j). In every case, the strongest cross-modal correlations fell within matching cortical territories, demonstrating that fUS-derived CBV changes are spatially aligned with optically measured hemoglobin dynamics and neuronal calcium activity.

Multitaper coherograms (9 tapers, 30 s window; Fig. 1k, Supp. Fig. 3) demonstrated strong coherence below 0.5 Hz between ΔCBV-fUS and hemoglobin changes, whereas ΔCa^2+^ coherence extended to higher frequencies, supporting the separate filter bands applied to each modality. Spatially averaged traces from the cortical ROIs (Fig. 1k) closely tracked one another across modalities, as quantified by z-scored joint distributions (Supp. Fig. 4a), demonstrating the sensitivity of fUS as its CBV signal closely matches optically measured hemoglobin dynamics. Across the cortex (Supp. Figs. 4b–e) and across mice (Fig. 1l, using region averaged traces), ΔCBV-fUS correlated positively with ΔHbO, ΔHbT, and ΔCa^2+^, and negatively with ΔHbR; neuronal peaks led ΔCBV-fUS by ~1.5 s (Fig. 1l), matching the expected neurovascular delay. Pearson’s r and Spearman’s ρ values were similar for every signal pair (Fig. 1m, Supp. Table 1), indicating an approximately linear relationship between fUS and optical measurements. Notably, the calcium correlation was weaker than the ΔHbT correlation (r = 0.34 vs. ~0.7). This is consistent with the fact that ΔCBV-fUS is driven by a combination of excitatory and inhibitory populations, of which the CaMKIIα-labeled excitatory neurons imaged here represent only one cell-type^7^.

Given the approximately linear relationship observed across modalities, we next tested whether simultaneous neuronal calcium and ΔCBV-fUS could be used to estimate a characteristic hemodynamic response function (HRF) during spontaneous brain activity^8,12^. Through least-squares deconvolution, we derived HRFs linking excitatory neuronal activity to hemoglobin and ΔCBV-fUS changes across distinct cortical regions (Supp. Fig. 4f) and across mice (Fig. 1n, using region averaged traces). These revealed a characteristic gamma-shaped transfer function with ~1.5 s time-to-peak for ΔCBV-fUS (Fig. 1o), and matching ΔHbT and ΔHbO HRFs; conversely, ΔHbR expectedly displayed a negative peak (predicted vs. measured traces in Supp. Fig. 4g, h).

While the above results demonstrate that neuronal activity is coupled to ΔCBV-fUS during spontaneous brain activity, underscoring the high baseline sensitivity of fUS to even unstimulated neural dynamics, we next tested whether this cross-modal relationship is preserved under evoked conditions with localized brain activation. Upon whisker stimulation, the strongest hemodynamic and neuronal activation (r > 0.6, p < 0.05) was localized to the somatosensory barrel cortex (Fig. 1p–r, Supp. Video 3). Coherence spectral maps, spatially averaged traces (Fig. 1s), and averaged cross-correlations (Supp. Fig. 5a) confirmed that ΔCBV-fUS correlates with hemoglobin and ΔCa^2+^ signals during stimulation, with neuronal peaks consistently preceding hemodynamic peaks by ~1.5 s. Trial-based HRFs (Fig. 1t) and predicted traces (Fig. 1u, Supp. Fig. 5b) showed strong agreement, confirming that cross-modal correspondence also holds during evoked activity.

To demonstrate the utility of simultaneous multimodal imaging in a complex brain disease setting, we conducted longitudinal fUS-optical imaging in an anesthetized orthotopic glioblastoma (GBM) mouse model at Days 9, 19, and 23 following GL261 tumor cell implantation (Supp. Fig. 6; Fig. 2a–c; Supp. Fig. 7a–c; Supp. Videos 4–6; Methods). Tumor-induced vascular disruption was independently confirmed via microbubble contrast-based ultrasound localization microscopy (Fig. 2d; Supp. Fig. 7d) utilizing the same imaging geometry outlined in Fig. 1a, and cross-validated by H&E staining (Fig. 2e; Supp. Fig. 7e) that closely matched the tumor boundaries observed on B-mode ultrasound (Supp. Fig. 6).

**Figure 2:**
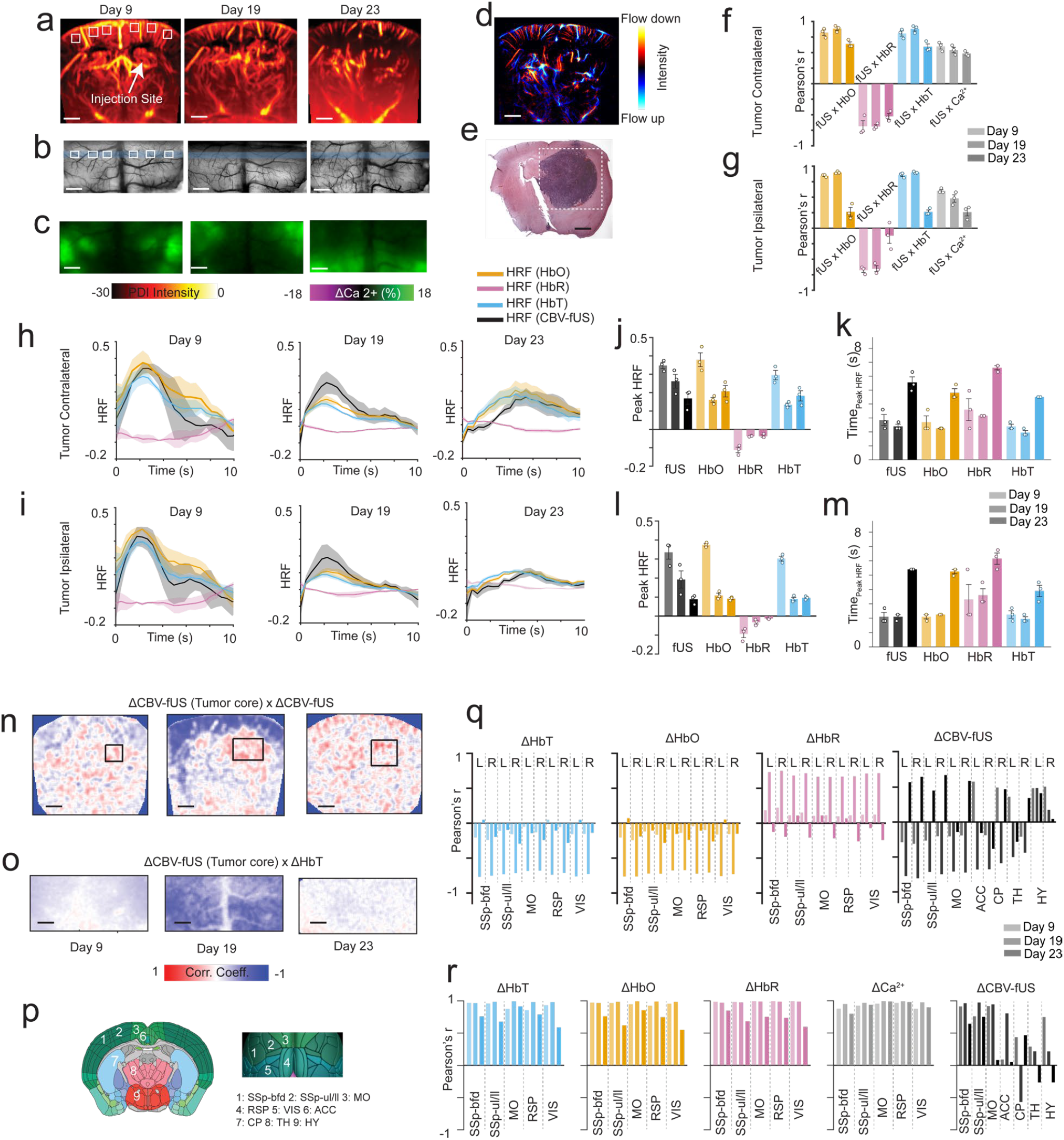
Simultaneous fUS-optical imaging for investigating glioblastoma neurovascular disruption. **(a)** fUS-derived cerebral blood volume (CBV) images, **(b)** optical reflectance (intrinsic optical signal imaging, IOSI) images, and **(c)** fluorescent neuronal calcium maps of an orthotopic brain tumor mouse at Days 9, 19, and 23 post-injection of GL261 glioblastoma cells 3 mm below the right cortex (injection site, arrow in (a)). **(d)** Ultrasound localization microscopy (ULM) image at Day 23. **(e)** H&E stain showing tumor extent correlates with in vivo B-mode ultrasound (Supp. Fig.6). **(f, g)** Pearson’s correlation between baseline-normalized CBV (ΔCBV-fUS) with baseline-normalized oxy-(ΔHbO), deoxy-(ΔHbR), and total hemoglobin (ΔHbT) and calcium (ΔCa^2+^) for contra- and ipsilateral cortex ROIs in (a, b). **(h, i)** Hemodynamic response functions (HRFs) in contra- and ipsilateral cortex across tumor progression. **(j, k)** HRF peak amplitude and time-to-peak, contralateral cortex; **(l, m)** same, ipsilateral. **(n, o)** Pixelwise correlation of averaged tumor-core ΔCBV-fUS (ROI inset in (n)) with ΔCBV-fUS and ΔHbT maps. **(p)** Allen mouse brain atlas regions used for analysis. **(q)** Regional correlation of ΔCBV-fUS with ΔHbO, ΔHbR, and ΔHbT across days. **(r)** Bilateral connectivity of ΔCBV-fUS, ΔHbO, ΔHbR, ΔHbT, and ΔCa^2+^ from regions in (q) across days. Scale bars, 1 mm. SSp-bfd, somatosensory barrel field; SSp-ul/ll, somatosensory limbs; MO, motor cortex; RSP, retrosplenial cortex; VIS, visual cortex; ACC, anterior cingulate cortex; CP, caudate putamen; TH, thalamus; HY, hypothalamus; PDI: Power Doppler Intensity.

We first asked whether GBM-induced disruption compromises the functional coupling between ΔCBV-fUS and optical signals. Simultaneous fUS-optical acquisition revealed that the correlation between ΔCBV-fUS and hemoglobin signals progressively declined in both ipsilateral and contralateral cortical regions (3 ROIs per side marked in Fig. 2a, b and Supp. Fig. 7a, b) with tumor progression (Fig. 2f, g; Supp. Fig. 7f, g). This reduced coupling could reflect a combination of local tissue necrosis, inflammation, glial scarring, or vascular remodeling, warranting further study. HRF analysis across fUS and optical hemodynamic signals at each timepoint revealed a progressive degradation of neurovascular coupling consistent with prior reports^13,14^, manifesting as reduced peak amplitude and delayed peak timing by Day 23 in both the contralateral (Fig. 2h– j; Supp. Fig. 7h–j) and ipsilateral (Fig. 2k–m; Supp. Fig. 7k–m) hemispheres (Supp. Fig. 8a,b). This bilateral impairment suggests that neurovascular uncoupling extends beyond the visible tumor boundary, a feature made apparent only through the wide spatial coverage of simultaneous fUS and optical imaging.

We then examined how hemodynamic activity within the tumor core relates to distal brain regions (Fig. 2n–o; Supp. Figs. 7n–o, 8c–d). At Day 9 and 19, the tumor core displayed inverse correlations with cortical ΔHbT, ΔHbO, and ΔCBV-fUS. By Day 23, these correlations reversed to positive for ΔCBV-fUS while remaining near zero for ΔHbT and ΔHbO, likely reflecting the superior depth sensitivity of fUS for capturing deep subcortical hemodynamic changes inaccessible to optical imaging. Region-wise correlation using the Allen mouse brain atlas^15^ (Fig. 2p, q; Supp. Fig. 7p) quantified these shifts. These findings are consistent with progressive coupling between tumor-associated vasculature and surrounding hemodynamics^16^. Bilateral connectivity analysis indicated potential progressive decoupling across cortical and subcortical structures for all hemodynamic signals by Day 23 (Fig. 2r; Supp. Fig. 7r). Interestingly, hemodynamic connectivity loss does not predict ΔCa^2+^ connectivity loss, suggesting that excitatory neuronal networks maintain partial functional integrity even as local vascular architecture severely deteriorates^17^. Overall, this highlights the potential of simultaneous fUS and optical imaging to study multiscale neurovascular dysfunction during disease progression.

In summary, we introduced a compact multimodal neuroimaging platform that enables simultaneous functional ultrasound, intrinsic optical signal, and widefield calcium imaging. In awake mice, these simultaneous measurements established robust spatial and temporal correspondence between fUS-derived CBV, optical hemoglobin signals, and neuronal calcium activity, while permitting the first direct cross-modal derivation of an HRF linking excitatory ΔCa^2+^ to ΔCBV-fUS during both spontaneous and sensory-evoked brain activity. Longitudinal glioblastoma imaging further revealed progressive neurovascular uncoupling, altered bilateral connectivity, and evolving relationships between superficial optical signals and deep hemodynamics that no single modality could resolve alone. Looking forward, the platform’s modular design will also serve as an open architecture to accommodate complementary modalities, such as photoacoustic molecular imaging of deep tissue^10^ (Supp. Fig. 8), B-mode ultrasound structural imaging (Supp. Fig. 6) and ultrasound localization microscopy (Figs. 2d, 7d), with combiner design files and analysis code provided to facilitate adoption. Additionally, as we measure calcium signals only from CaMKIIα-positive excitatory neurons, future implementations with inhibitory, astrocytic, or vascular reporters will be needed to decompose cell-type-specific contributions to fUS-derived CBV. More broadly, by simultaneously capturing CBV, molecular hemodynamics, and neuronal activity, the fUS-optical framework will facilitate mechanistic interpretation of the fUS signal—an understanding of increasing importance as functional ultrasound expands into clinical neuroimaging and brain-computer interface applications.

## Methods

### Animal Preparation

#### Cranial window and head-bar implantation

Transgenic mice (3-6 months old, n=4 for awake head restraint study, n=2 for brain cancer study under low dose (0.5 – 1%) anesthesia, male or female) were used for this study and generated by crossing CaMKII-Cre (#005359, Jackson Labs) with LSL-jGCaMP8s (#037952, Jackson Labs) lines. These survival surgical procedures were followed as previously described^8,18^. In brief, mice were anaesthetized using isoflurane (3-5%) and medical oxygen supply mixture (1 L/min) and transferred to a stereotaxic frame with continuous supply of anesthetic gas (1-3% Isoflurane mixed with 1 L/min oxygen). A warming pad was placed below the mice and a rectal thermal probe was inserted to maintain mice’s body temperature at 37°C. Hair was removed using a depilatory cream and the exposed skin was cleaned by alternating between betadine and 70% ethanol three times. Bupivacaine was injected subcutaneously as a local anesthetic. Following this, a midline incision was made and connective tissue cleared to expose the skull surface. For the two GBM mice, 250,000 GL261 murine glioma cells loaded in 5 μL of phosphate buffer saline were injected through a 0.5 mm drilled burr hole (Coordinates: anteroposterior (AP) −1.0 mm, mediolateral (ML) −2.0 mm, and dorsoventral (DV) −2.0 mm). The craniotomy was done next by drilling over a rectangular window ~4×8 mm^2^ (AP +0.5 to AP −3.5 mm) using 0.5 mm dental drill bit (19007-05, Fine Science Tools Inc., California, USA). Care was taken to regularly cool down the drilled area with sterile saline. Drilling was done until the skull felt like a “floating island”. The skull was then slowly lifted using fine forceps carefully avoiding any bleeds and the dura was left intact. A 50 μm thick PMP film (MX002, Goodfellow Corp, Utah, USA) trimmed to the size of the cranial window was placed over the dura. The film was sealed using cyanoacrylate glue and dental cement (C&B Metabond, Parkell, NY, USA). For awake mice imaging, a head bar was additionally attached over the interparietal bone with cyanoacrylate glue. A layer of dental cement and another layer of cyanoacrylate glue was added around the edges and over the fixed head bar to further reinforce and seal the cranial window. The mice were administered post-operative analgesia (Buprenorphine Ethiqa XR) subcutaneously to aid in their recovery and minimize discomfort. The mice were given seven days to recover followed by three days of incremental habituation over the head-fixing apparatus in case of awake imaging. All animal care and procedures were carried out in accordance with the Institutional Animal Care and Use Committee (IACUC) at The Pennsylvania State University.

#### fUS-Optical Imaging via Custom, Compact Acousto-Optic Imaging Head

fUS-optical multimodal imaging was acquired with a custom-designed, 3D-printed acousto-optic combiner that served as the fUS-optical imaging head (Figure 1(a)–(c), Supp. Fig. 1). The imaging head consists of a 45°-oriented cover glass slip (Cat# 480393-092, VWR International, Pennsylvania, USA) surrounded by degassed 1% agarose gel (Cat# 20-102GP, Apex, Genesee Scientific, California, USA) medium. The cover glass reflected the plane wave ultrasound beam onto the optical axis, and the agarose provided acoustic coupling. The optical characterization of the imaging head was done using USAF 1951 resolution target and Rayleigh’s 26.3% intensity criteria was used for obtaining spatial resolution (Supp. Fig. 2). The ultrasound transducer probe (Vermon, Tours, France, 14.4 MHz, 40% fractional bandwidth, 128 elements with 0.1 mm pitch, 1.5 mm elevation, and 6 mm elevational focus) was fixed laterally inside the imaging head, pressing against the solidified agarose medium with a thin layer of ultrasound gel (Sterile Aquasonic 100, Parker Laboratories, New Jersey, USA). The ultrasound probe was fixed in place using a magnetic cover. A second cover glass slip was placed on top of the agarose with a thin layer of water below for optical refractive index matching and improving optical transparency. The fUS-optical imaging head was further coupled to the mice PMP cranial window with a thin layer of ultrasound gel to ensure good acoustic coupling. Care was taken to avoid any bubbles in the ultrasound gel and any residual bubbles were removed using a micropipette. Awake fUS-optical imaging of healthy mice was performed after acclimation to head restraint. Longitudinal fUS-optical imaging of GBM mice was performed under low-dose (0.5–1%) isoflurane anesthesia.

For fUS imaging, blocks of 200 compounded frames at 500 Hz compound frame rate (11 plane waves, angled from −15° to 15°) were collected at a voltage 20.3 V using the ultrasound transducer probe connected to a multichannel ultrasound data acquisition system (Vantage 256 system, Verasonics Inc., Washington, USA). In awake mice, transmit apodization was applied by excluding 24 to 32 elements on each side of the aperture with a Hanning window to reduce motion artifacts from facial muscle movements. fUS imaging provides a spatial resolution of ~100 μm × 100 μm (width × depth), with an imaging plane thickness of approximately ~400 μm^19^.

For optical imaging (IOSI and widefield fluorescence), 470 nm, 530 nm, and 625 nm LEDs (M470L5, M530L4, M625L4, Thorlabs, New Jersey, USA) illuminated the mice brain through the optical cavity of the imaging head. The three wavelengths were interleaved at 20 Hz frame rate with illumination timing synced to CMOS camera (IDS U3-3040CP-M-GL, IDS Imaging Development Systems Inc., Massachusetts, USA) acquisition. A long pass 515 nm optical filter (AT515LP, Chroma Technology Corp., Vermont, USA) was placed in between the objective lens (Nikon AF Micro Nikkor 60 mm f/2.8D lens, Nikon, Tokyo, Japan) and the CMOS camera to block 470 nm GCaMP fluorescence excitation wavelength. The camera exposure and LED illumination time was matched to 45 ms. Whisker stimulation was provided using a custom engineered relay-based air-puff system. The data acquisition software was written using a MATLAB code and timing was synchronized with Arduino Uno R3 (Arduino, Monza, Italy). The fUS and optical imaging modalities ran asynchronously and were matched using timestamps thereby ensuring proper synchronization. For co-registration of the imaging modalities, we used the tip of the window as the fiducial marker and the translation of the imaging head was noted to identify the fUS imaging plane on the cortex.

For ULM imaging, 400 frames at 1000 Hz compound frame rate (5 plane waves angle at −15° to 15°) were collected at a voltage 5.3 V following 100 μL bolus retro-orbital injection of custom-made microbubbles.

For combined fUS and photoacoustic imaging, we used the methodology in^10^ with fUS images acquired through 150 compounded frame at 1200 Hz compounded frame rate (11 plane waves angle at −7° to 7°) with photoacoustic image frames captured at 800 nm (Phocus MOBILE, Opotek, California, USA).

#### Ultrasound Data Processing

Using ultrafast ultrasound approach, plane-wave acquisitions at different tilt angles were beamformed and coherently compounded. The compounded frame blocks were filtered using singular value decomposition (SVD) spatio-temporal filter^20^ by discarding top singular values (60 for awake mice, 30 for anaesthetized brain cancer mice). Each compounded frame block was further high pass filtered with 20 Hz frequency cutoff to generate Power Doppler Intensity (PDI) map proportional to CBV. The two filters ensured that CBV signals can be separated from tissue motion. Subsequently, pixel wise change in CBV maps were generated to derive baseline normalized ΔCBV-fUS using the formula:

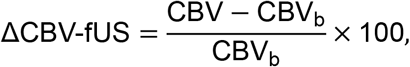

where CAV_b_ is the mean intensity over the baseline (15 s for non-stimulated data set and 5 s for stimulus evoked activity). B-mode ultrasound images were extracted from the ultrasound beamformed data without filtering. ULM maps were constructed using the codes and methodology adapted from^21^. In brief, we used an SVD filter of 12, 100 particles per frame, localized to sub-pixel resolution using spline-based interpolation. Thereafter, tracks were formed using the Hungarian algorithm^22^ and were subsequently accumulated for forming ULM maps.

#### Optical Data Processing

Intrinsic optical signal image (IOSI) analysis: IOSI signals rely on reflectance changes of the illuminating wavelengths. Here, 530 nm and 625 nm were used to unmix HbO and HbR signals using the modified Beer-lambert’s law methodology mentioned in ^23^. Briefly, the relative photon intensity at wavelength (λ) travelling through tissue is given as

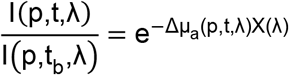

where p is the pixel location, t is the time, t_b_ is the baseline (15 s for non-stimulated data set and 5 s for stimulus evoked activity), Δµ_a_(p, t, λ) is the change in absorption coefficient at time t relative to baseline and X(λ) is the wavelength dependent differential path length factor. Moreover, Δµ_a_(p, t, λ) depends on the molar extinction coefficients ϵ_n_(λ) and relative chromophore concentrations (ΔC_n_(p, λ)). Δµ_a_(p, t, λ) is therefore given as:

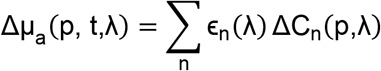

For the chosen wavelengths of 530 nm and 625 nm, the dominant absorbers are oxy- and deoxy-hemoglobins. By using the two equations above, we calculated the relative ΔHbO and ΔHbR corresponding to each pixel and time.

Widefield Ca^2+^ image analysis: The emitted fluorescence wavelength from the GCaMP fluorophore is contaminated by hemodynamic changes. This was corrected by dividing the fluorescence ratio by the reflectance ratio (relative to baseline) of 530 nm reflectance maps, as mentioned in ^23^. Following this, changes in neuronal calcium signals ΔCa^2+^ were calculated

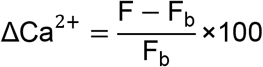

Where F_b_ is mean intensity over the baseline (15 s for non-stimulated data set and 5 s for stimulus evoked activity).

#### Correlation and Coherence Analysis

Optical signals were downsampled to the functional ultrasound frame rate prior to spectral and correlation analysis. Spectral coherence maps were computed for unmixed hemoglobin concentrations (ΔHbO, ΔHbR and ΔHbT), neuronal calcium (ΔCa^2+^) and cerebral blood volume (ΔCBV-fUS) using the multitaper method implemented in the Chronux toolbox^24^. A time– bandwidth product of 5 with 9 Slepian tapers was applied within a 30 s sliding window advanced in 2 s steps, restricting the analysis to the 0–1 Hz frequency band. Correlation analysis was performed after bandpass filtering the hemodynamic signals (ΔCBV-fUS, ΔHbO, ΔHbR and ΔHbT) to 0.02–0.5 Hz for awake mice and 0.02–0.2 Hz for anaesthetized glioblastoma mice, removing slow drifts and cardiac frequency contributions. The change in upper limit of filtering bands from 0.5 Hz (awake mice) to 0.2 Hz (anesthesia mice) was to account for anaesthesia-induced slowing of vascular dynamics. Calcium signals (ΔCa^2+^) were high pass filtered with a cutoff frequency of 0.02 Hz to preserve slow hemodynamic fluctuations while retaining fast neural responses. The relationship between z-scored ΔCBV-fUS and each optical signal was assessed by plotting their joint distribution and estimating the slope via linear regression. Calcium signals were temporally shifted by ~1.5 s to account for the neurovascular coupling lag prior to correlation analysis. We used the Matlab xcorr function to obtain correlation traces and the corr function to obtain Pearson’s r and Spearman’s ρ coefficient. To generate the ROI-seed cross correlation maps, the change maps (ΔCBV-fUS, ΔHbT and ΔCa^2+^) were spatially averaged within the ROI and pixel by pixel cross-correlated cross modality to generate the correlation maps. For tumor core connectivity analysis, the spatially averaged ΔCBV-fUS signal extracted from the tumor core was correlated pixel-wise with ΔCBV-fUS, ΔHbT, ΔHbO and ΔHbR maps within a ±1 s lag window to generate the correlation map. For region specific analysis ROIs were manually delineated according to a standard brain atlas, and peak connectivity between each ROI and the tumor core was determined within a ±1 s lag window. The lag window was used to account for hemodynamic disruption that may delay hemodynamic changes. Bilateral connectivity was assessed by comparing signal pairs from left hemisphere (contralateral) and right hemisphere (ipsilateral) counterparts with zero lag correlation.

#### Hemodynamic Response Function

The filtered signals, same as for correlation analysis, were also used for HRF analysis. To estimate the HRF from ΔCa^2+^ signals, we used a deconvolution approach formulated as a least squares problem with Tikhonov regularization (λ=0.01) to ensure a stable solution ^25^. Briefly, we expressed the neurovascular response as a linear system

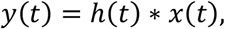

where y(t) is the output hemodynamic signal (ΔCBV-fUS, ΔHbT, ΔHbR and ΔHbO), x(t) is the input ΔCa^2+^ neuronal response, and h is the HRF to be estimated. In matrix-vector form, this is expressed as:

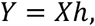

where X is a Toeplitz convolution matrix derived from the neuronal signal. The HRF (h) was estimated by minimizing the following objective function:

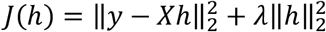

where λ is the regularization parameter. The closed-form solution to the minimization problem above is given as:

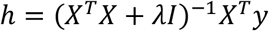

This was computed iteratively using the MATLAB lsqr function. We assumed the length of the HRF to be 9 s (20 samples at the fUS frame rate) for awake mice data and 13.6 s (30 samples at the fUS frame rate) for the anaesthetized brain cancer imaging data to account for anaesthesia-induced slowing of neurovascular dynamics. The HRFs were calculated over the spatially averaged traces. HRFs were computed using 100 s signal trace for non-stimulated awake and anaesthetized mice data and 25 s for whisker stimulation data (5s pre baseline, 5s post baseline).

#### Histology

The mice were euthanized on day 24 post GL261 injection. The skull with brain intact was immersed in 4% paraformaldehyde (PFA) in 0.1 M phosphate-buffered saline (PBS, pH 7.4) for fixation. After fixation, the specimens were transferred to Decalcifying Solution B for decalcification, after which the brain tissue was extracted. The specimens were then dehydrated through a graded ethanol series, cleared in xylene, and infiltrated with paraffin wax. Tissues were embedded in paraffin and sectioned at 10 µm. Sections were mounted onto glass slides and stained with hematoxylin and eosin. Stained sections were cover slipped and imaged using a custom light microscope.

#### Statistics

We report both Pearson’s r and Spearman’s ρ for assessing correlation, allowing us to quantify relationships under both normal and non-normal distribution assumptions between ΔCBV-fUS and optical signals. Statistical significance of correlations was assessed using a two-tailed t-test against zero, with a threshold of p < 0.05. For group-level comparisons across signal types, paired t-tests against zero with Bonferroni correction for multiple comparisons were used. *p<0.05, **p<0.01, ***p<0.001.

## Acknowledgement

We thank Richelle Irene Brown (The Pennsylvania State University) and Animal diagnostic laboratory, The Pennsylvania State University for their help preparing histological samples.

## Contributions

**Shubham Mirg**: Conceptualization, Methodology, Software, Data curation, Visualization, and Investigation; **Prameth Gaddale**: Investigation; **Achutha Kumar**: Investigation and Visualization; **Krishnendu Samanta**: Investigation; **Bhawna Saini**: Investigation; **Saniya P. Patil**: Investigation; **Ana A. Vargas**: Resources; **Aayushi Laliwala**: Resources; **Agata A. Exner**: Resources; **Yong Wang**: Resources; **Grayson O. Sipe**: Methodology and Resources; **Sri-Rajasekhar Kothapalli**: Conceptualization, Data curation, Methodology, Resources, Investigation and Supervision. Shubham Mirg wrote the original draft, and the co-authors provided comments and edited the manuscript.

