## Supplementary Text for "Simultaneous Functional Ultrasound, Intrinsic Optical Signal and Widefield Calcium Neuroimaging"

**Supplementary Table 1: Comparison of Pearson and Spearman correlation coefficients for the correlation between ΔCBV-fUS and optical signals across mice (n=4).**

**Supplementary Figure 1: 3D printed fUS-optical imaging head**

**Supplementary Figure 2: Optical characterization of fUS-optical imaging setup**

**Supplementary Figure 3: Coherence relationship between different signals obtained using fUS-optical imaging**

**Supplementary Figure 4: Regional variability analysis for correspondence between cortical fUS and optical signals.**

**Supplementary Figure 5: Cross correlation and predicted trace correlation for whisker stimulation**

**Supplementary Figure 6: In vivo Ultrasound B-mode images of orthotopic glioblastoma tumor progression**

**Supplementary Figure 7: Additional data for fUS-optical imaging of glioblastoma mouse**

**Supplementary Figure 8: Longitudinal study of predicted-trace correlation and tumor core connectivity during tumor growth**

**Supplementary Figure 9: In vivo functional ultrasound and photoacoustic imaging of mouse brain**

**Supplementary Video 1: fUS-optical awake mice imaging**

**Supplementary Video 2: fUS-optical change maps of spontaneous awake mice brain activity**

**Supplementary Video 3: fUS-optical change maps of awake mice brain activity during whisker stimulus**

**Supplementary Video 4: fUS-optical change maps of spontaneous brain activity in anaesthetized glioblastoma mice at Day 9**.

**Supplementary Video 5: fUS-optical change maps of spontaneous brain activity in anaesthetized glioblastoma mice at Day 19**

**Supplementary Video 6: fUS-optical change maps of spontaneous brain activity in anaesthetized glioblastoma mice at Day 23**

**Supplementary Table 1: Comparison of Pearson and Spearman correlation coefficients for the correlation between ΔCBV-fUS and optical signals across mice (n=4).**

|  | Pearsons’s r | p value | Spearman’s ρ | p value |
| --- | --- | --- | --- | --- |
| ΔCBV-fUS x ΔHbO | 0.719 ± 0.043 | 0.0009 | 0.712 ± 0.035 | 0.0005 |
| ΔCBV-fUS x ΔHbR | -0.653 ± 0.042 | 0.0009 | -0.623 ± 0.068 | 0.0042 |
| ΔCBV-fUS x ΔHbT | 0.681 ± 0.076 | 0.0048 | 0.680 ± 0.047 | 0.0015 |
| ΔCBV-fUS x ΔCa^2+^ | 0.335 ± 0.110 | 0.0433 | 0.350 ± 0.072 | 0.0119 |

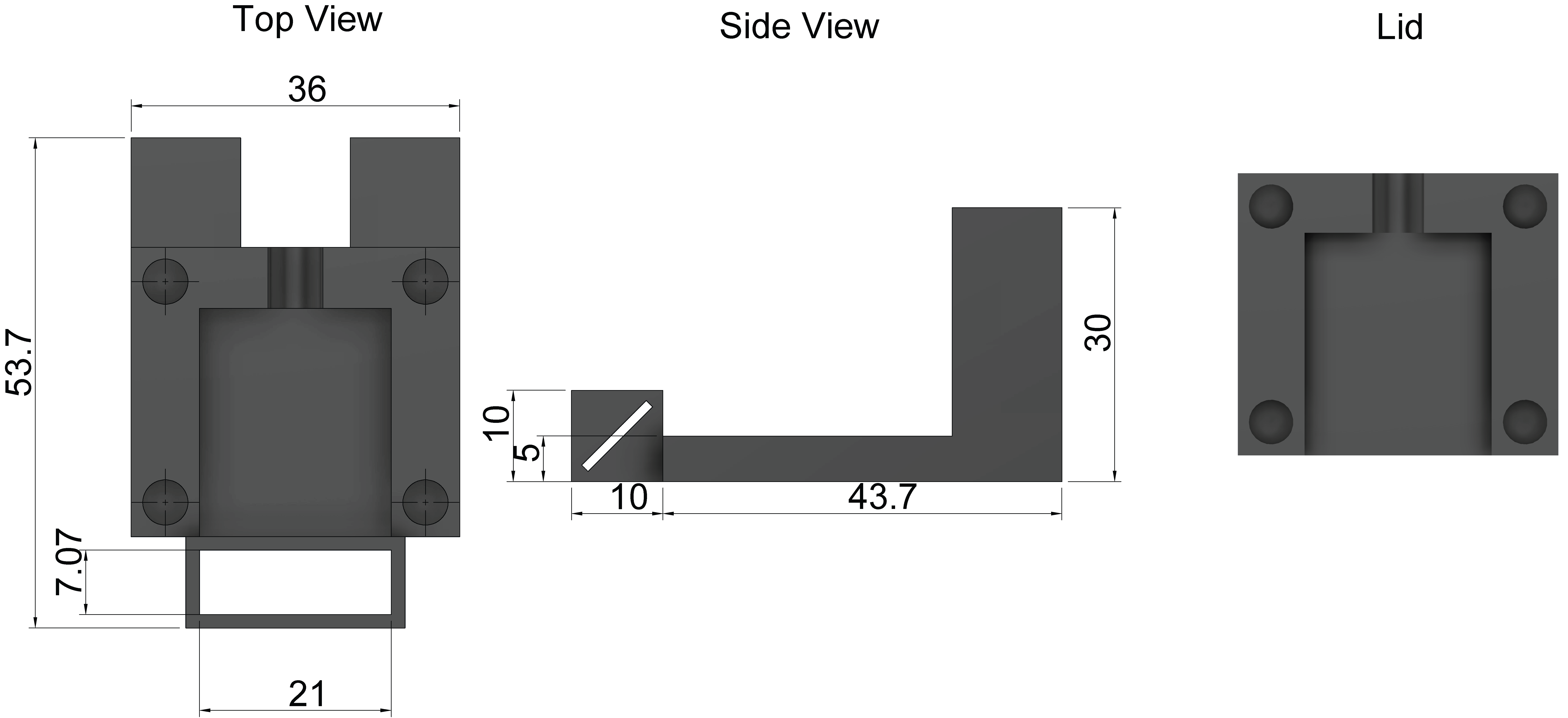

**Supplementary Figure 1: 3D printed fUS-optical imaging head**

Dimensions of the custom designed 3D printed fUS-optical imaging head with a 45° cover glass that allows for placement of ultrasound transducer horizontally thereby allowing for combing fUS with optical imaging. All dimensions in mm.

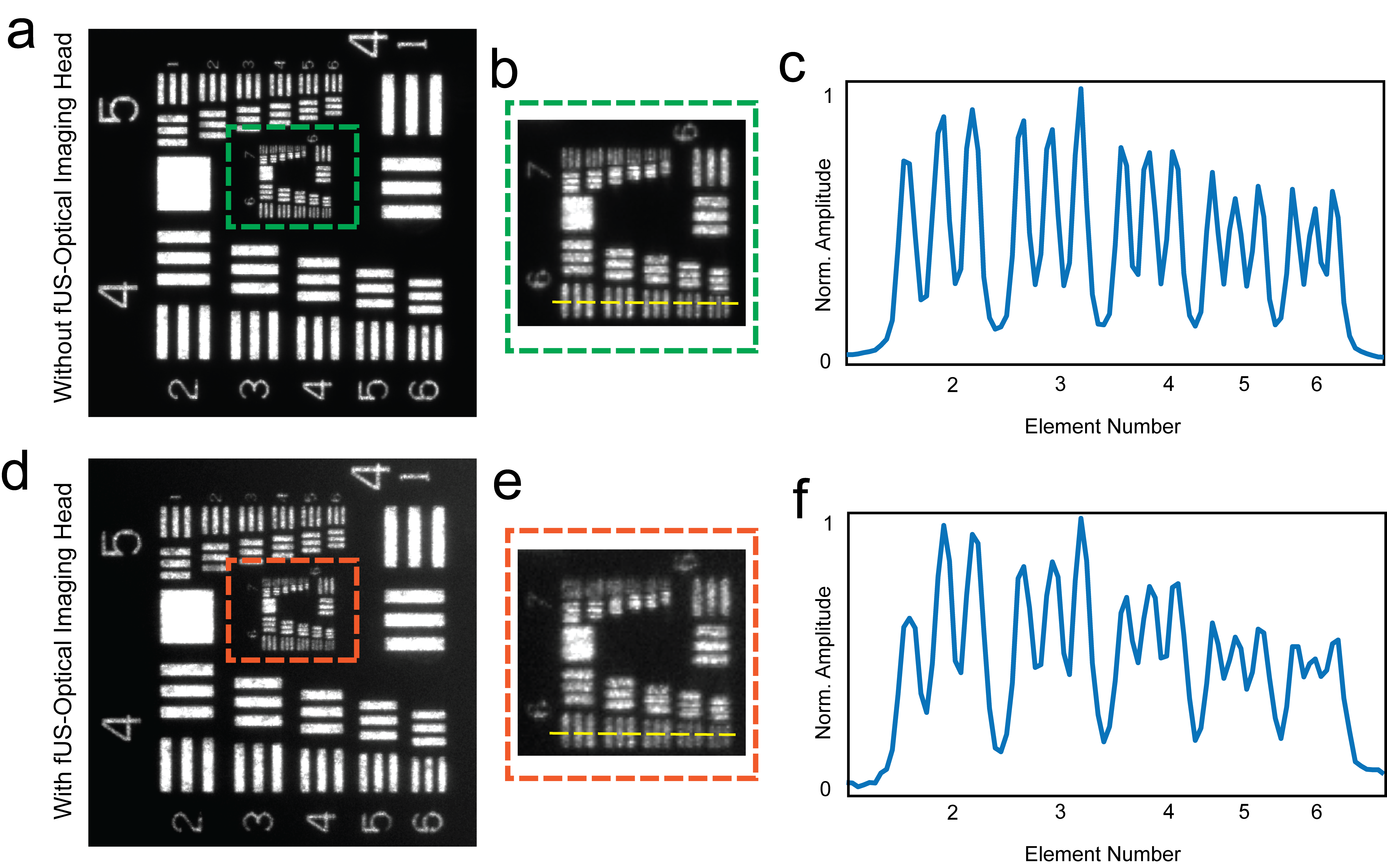

**Supplementary Figure 2: Optical characterization of fUS-optical imaging setup**

**(a)** Optical image of USAF 1951 resolution target without fUS-optical imaging head. **(b)** Green insert magnified image of USAF 1951 resolution target. **(c)** Intensity across the line marked in (b). **(d)** Optical image of USAF 1951 resolution target with fUS-Optical imaging head. **(e)** Orange insert magnified image of USAF 1951 resolution target. **(f)** Intensity across the line marked in (e).

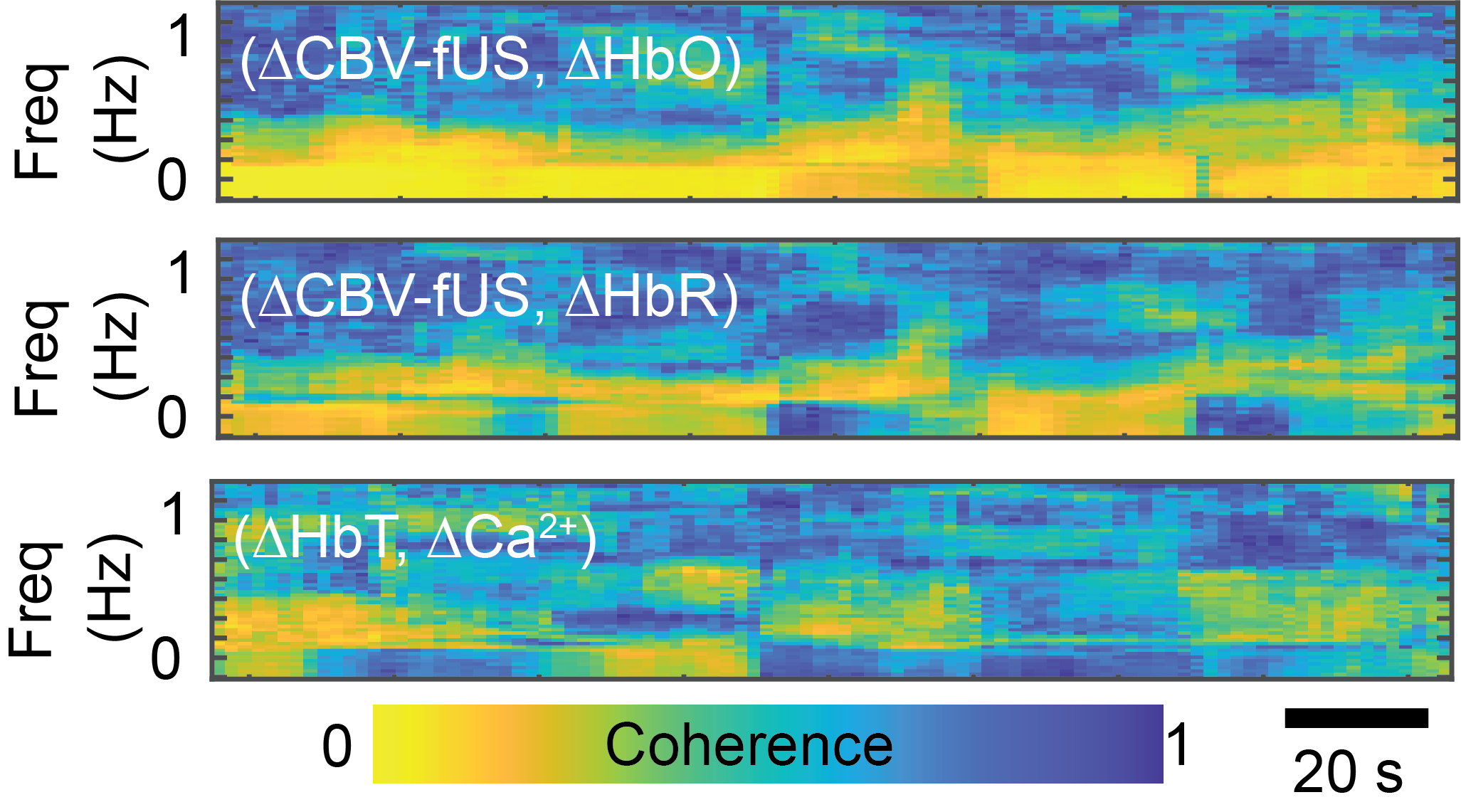

**Supplementary Figure 3: Coherence relationship between different signals obtained using fUS-optical imaging**

Spectral multitaper coherogram of baseline-normalized cerebral blood volume (ΔCBV-fUS) with oxy- (ΔHbO) and deoxy-hemoglobin (ΔHbR), with the coherogram map between total hemoglobin (ΔHbT) and calcium (ΔCa²⁺) shown at the bottom.

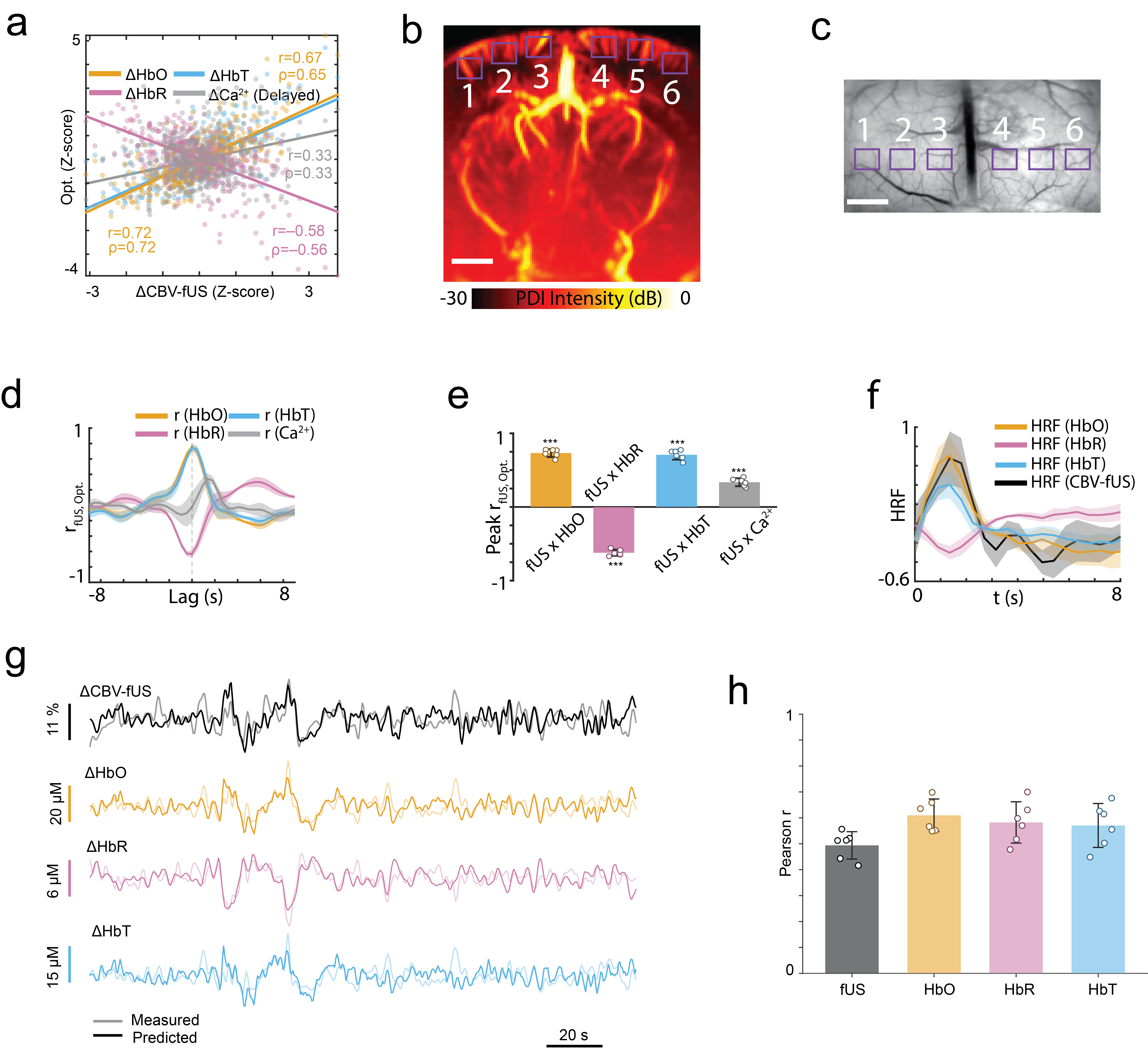

**Supplementary Figure 4: Regional variability analysis for correspondence between cortical fUS and optical signals.**

**(a)** Comparison of z-scored baseline-normalized cerebral blood volume (ΔCBV-fUS) with oxy- (ΔHbO), deoxy-hemoglobin (ΔHbR), and calcium (ΔCa²⁺) over the ROI in Fig. 1e, f. **(b)** fUS and **(c)** optical regions of interest (ROIs) for analysis across cortex. **(d)** Cross-correlation of ΔCBV-fUS with ΔHbO, ΔHbR, total hemoglobin (ΔHbT), and ΔCa²⁺ over the six ROIs in (b, c). (e) Corresponding correlation peaks from (d). **(f)** Hemodynamic response functions (HRFs) linking calcium changes to ΔCBV-fUS and intrinsic optical signal imaging (IOSI)-derived hemoglobin signals (6 ROIs). **(g)** HRF-predicted traces overlaid on measured ΔCBV-fUS, ΔHbO, ΔHbR, and ΔHbT traces for ROI 6 in (b, c). **(h)** Pearson’s correlations between predicted and measured ΔCBV-fUS, ΔHbO, ΔHbR, and ΔHbT traces for the 6 ROIs. Scale bars, 1 mm.

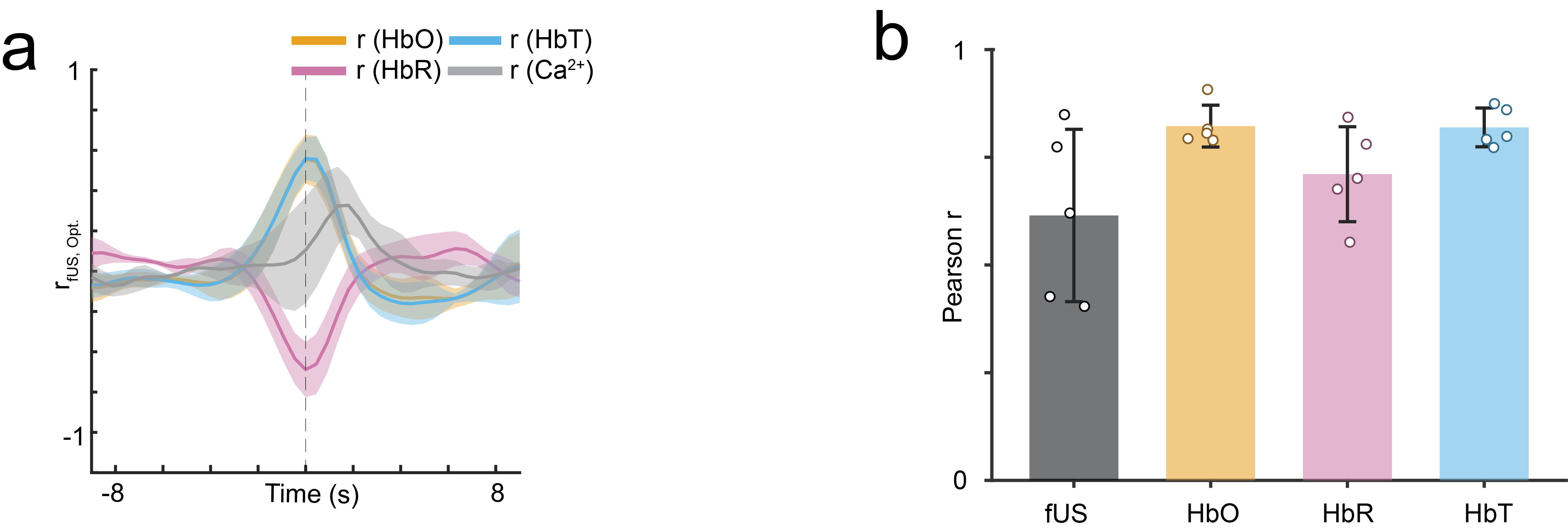

**Supplementary Figure 5: Cross correlation and predicted trace correlation for whisker stimulation**

**(a)** Cross-correlation of baseline-normalized cerebral blood volume (ΔCBV-fUS) with oxy- (ΔHbO), deoxy- (ΔHbR), total hemoglobin (ΔHbT), and calcium (ΔCa²⁺) signals for a 15 s whisker stimulus (n = 3, 5 whisker stimuli per mouse). **(b)** Predicted-trace correlations with measured ΔCBV-fUS, ΔHbO, ΔHbR, and ΔHbT obtained using a ΔCa²⁺-linked hemodynamic response function (HRF) for the whisker-stimulus traces in Fig. 1u.

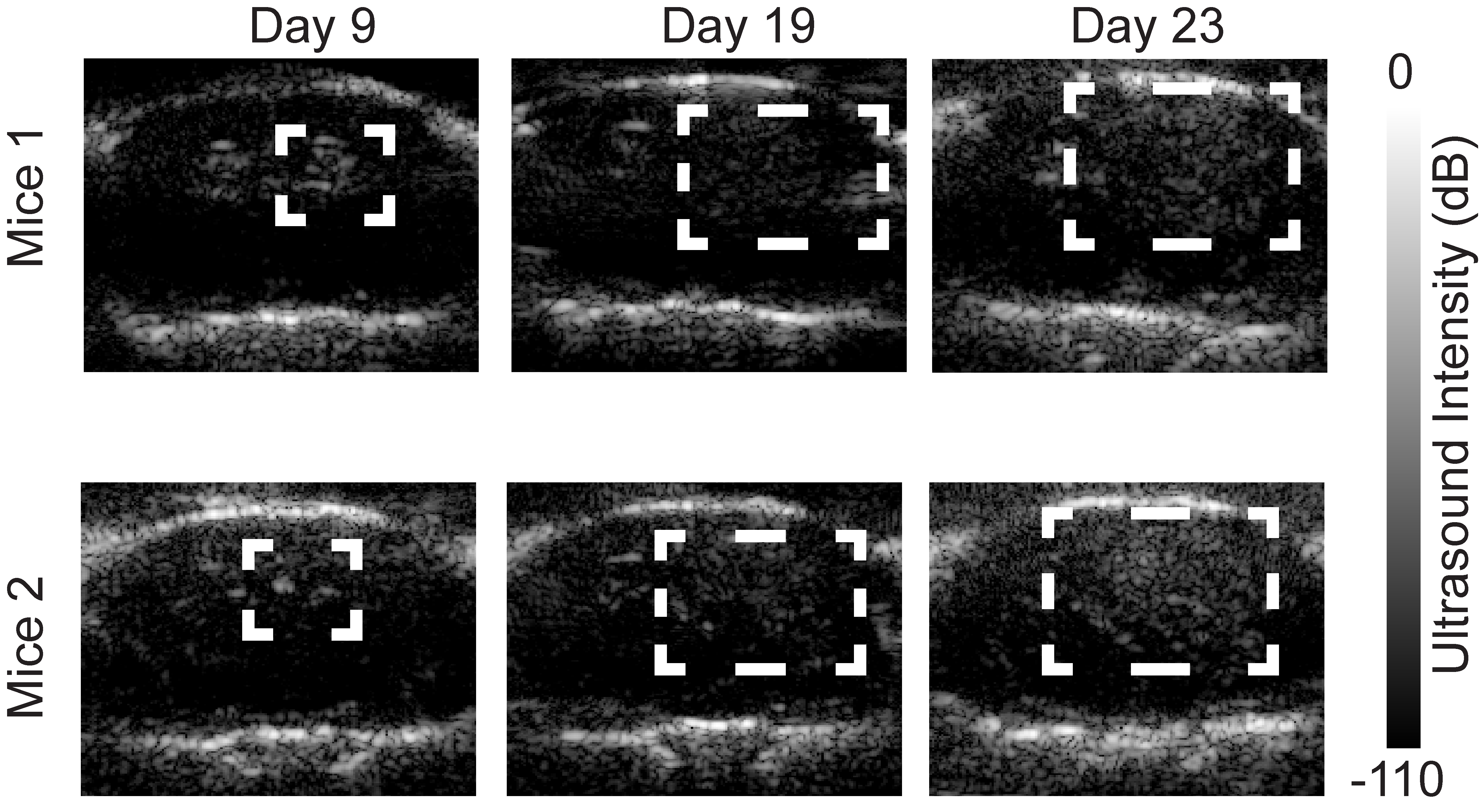

**Supplementary Figure 6: In vivo Ultrasound B-mode images of orthotopic glioblastoma tumor progression**

The panel shows in vivo ultrasound B-mode images of orthotopic glioblastoma tumor progression over 23 days for mice 1 and 2, obtained using fUS-optical imaging setup. The white inset indicates the progression of tumor made apparent in structural ultrasound images by stiffness (mechanical impedance) changes of tumor core.

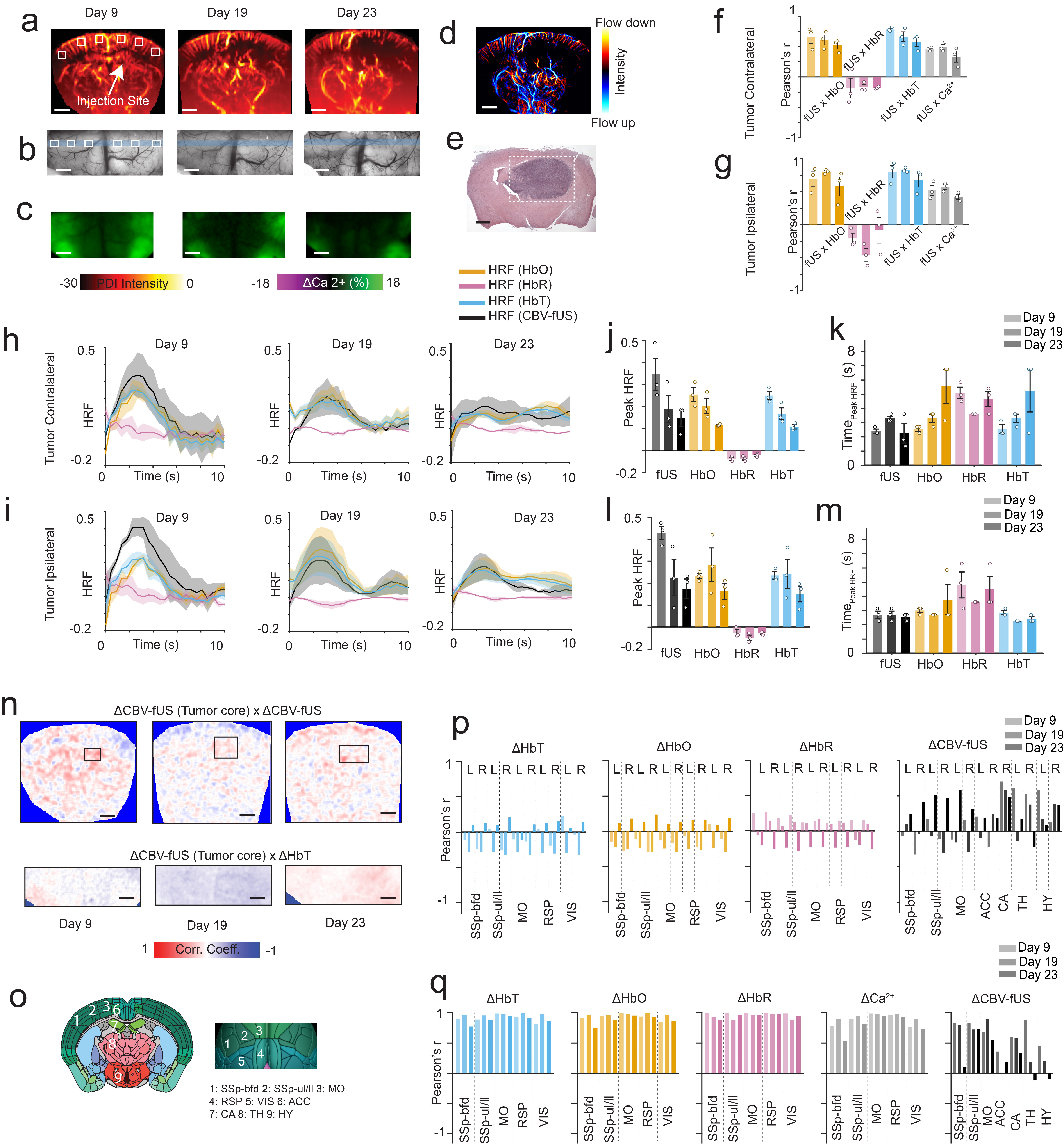

**Supplementary Figure 7: Additional data for fUS-optical imaging of glioblastoma mouse**

**(a)** fUS-derived cerebral blood volume (CBV) images, **(b)** optical reflectance (intrinsic optical signal imaging, IOSI) images, and **(c)** fluorescent neuronal calcium maps of an orthotopic brain tumor mouse at Days 9, 19, and 23 post-injection of GL261 glioblastoma cells 3 mm below the right cortex (injection site, arrow in (a)). **(d)** Ultrasound localization microscopy (ULM) image at Day 23. **(e)** H&E stain showing tumor extent correlates with in vivo B-mode ultrasound (Supp. Fig.6). **(f, g)** Pearson’s correlation between baseline-normalized CBV (ΔCBV-fUS) with baseline-normalized oxy- (ΔHbO), deoxy- (ΔHbR), and total hemoglobin (ΔHbT) and calcium (ΔCa²⁺) for contra- and ipsilateral cortex ROIs in (a, b). **(h, i)** Hemodynamic response functions (HRFs) in contra- and ipsilateral cortex across tumor progression. **(j, k)** HRF peak amplitude and time-to-peak, contralateral cortex; **(l, m)** same, ipsilateral. **(n, o)** Pixelwise correlation of averaged tumor-core ΔCBV-fUS (ROI inset in (n)) with ΔCBV-fUS and ΔHbT maps. **(p)** Allen mouse brain atlas regions used for analysis. **(q)** Regional correlation of ΔCBV-fUS with ΔHbO, ΔHbR, and ΔHbT across days. **(r)** Bilateral connectivity of ΔCBV-fUS, ΔHbO, ΔHbR, ΔHbT, and ΔCa²⁺ from regions in (q) across days. Scale bars, 1 mm. SSp-bfd, somatosensory barrel field; SSp-ul/ll, somatosensory limbs; MO, motor cortex; RSP, retrosplenial cortex; VIS, visual cortex; ACC, anterior cingulate cortex; CA, Hippocampus; TH, thalamus; HY, hypothalamus; PDI: Power Doppler Intensity.

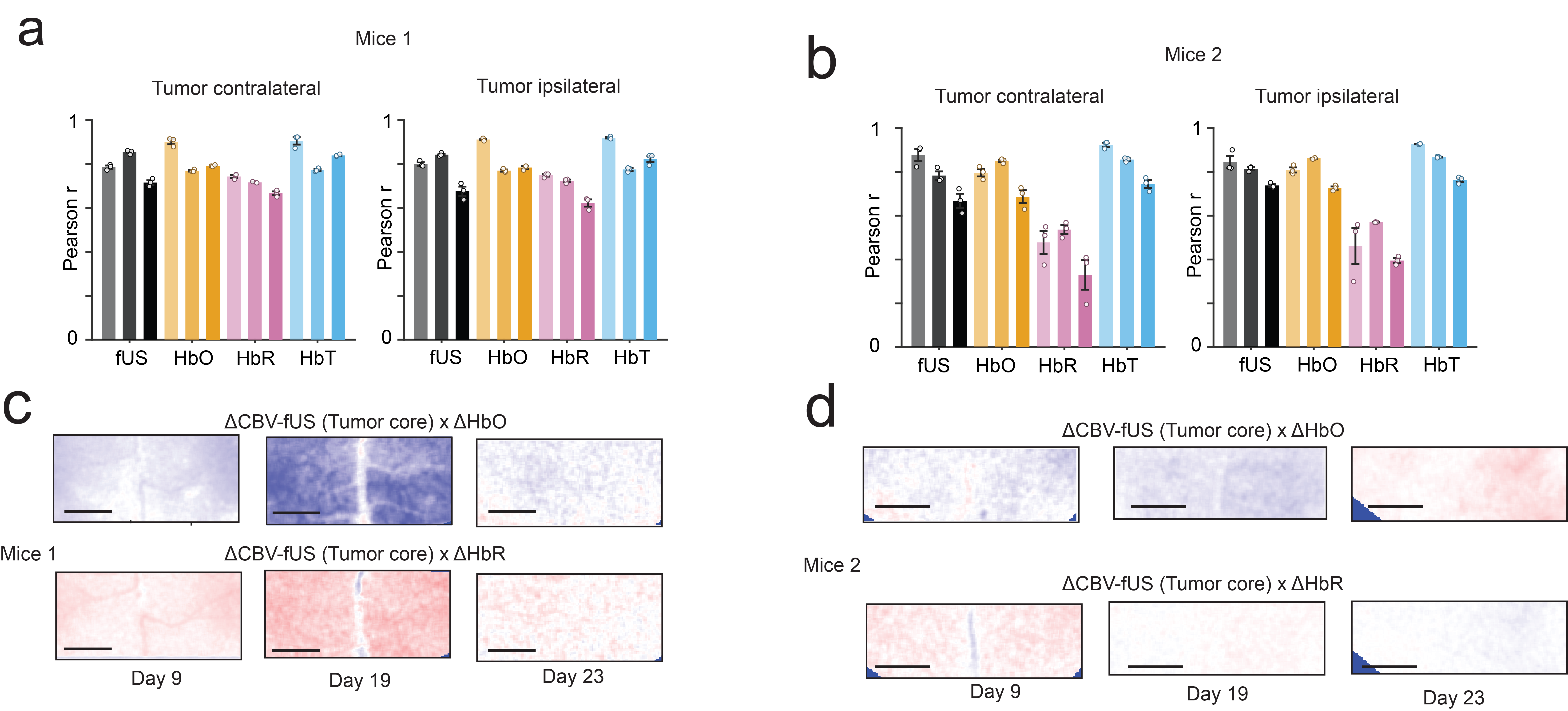

**Supplementary Figure 8: Longitudinal study of predicted-trace correlation and tumor core connectivity during tumor growth**

**(a)** Predicted-trace correlations with measured baseline-normalized cerebral blood volume (ΔCBV-fUS), oxy- (ΔHbO), deoxy- (ΔHbR), and total hemoglobin (ΔHbT), obtained from a ΔCa²⁺-linked hemodynamic response function (HRF) across days for Mouse 1. **(b)** Same as (a) for Mouse 2. **(c)** Pixelwise correlation of spatially averaged tumor-core ΔCBV-fUS (ROI inset in Fig. 2n) with ΔCBV-fUS, ΔHbO, and ΔHbR maps. **(d)** Same as (c) for Mouse 2 (ROI inset in Supp. Fig. 7n). Scale bars, 2 mm.

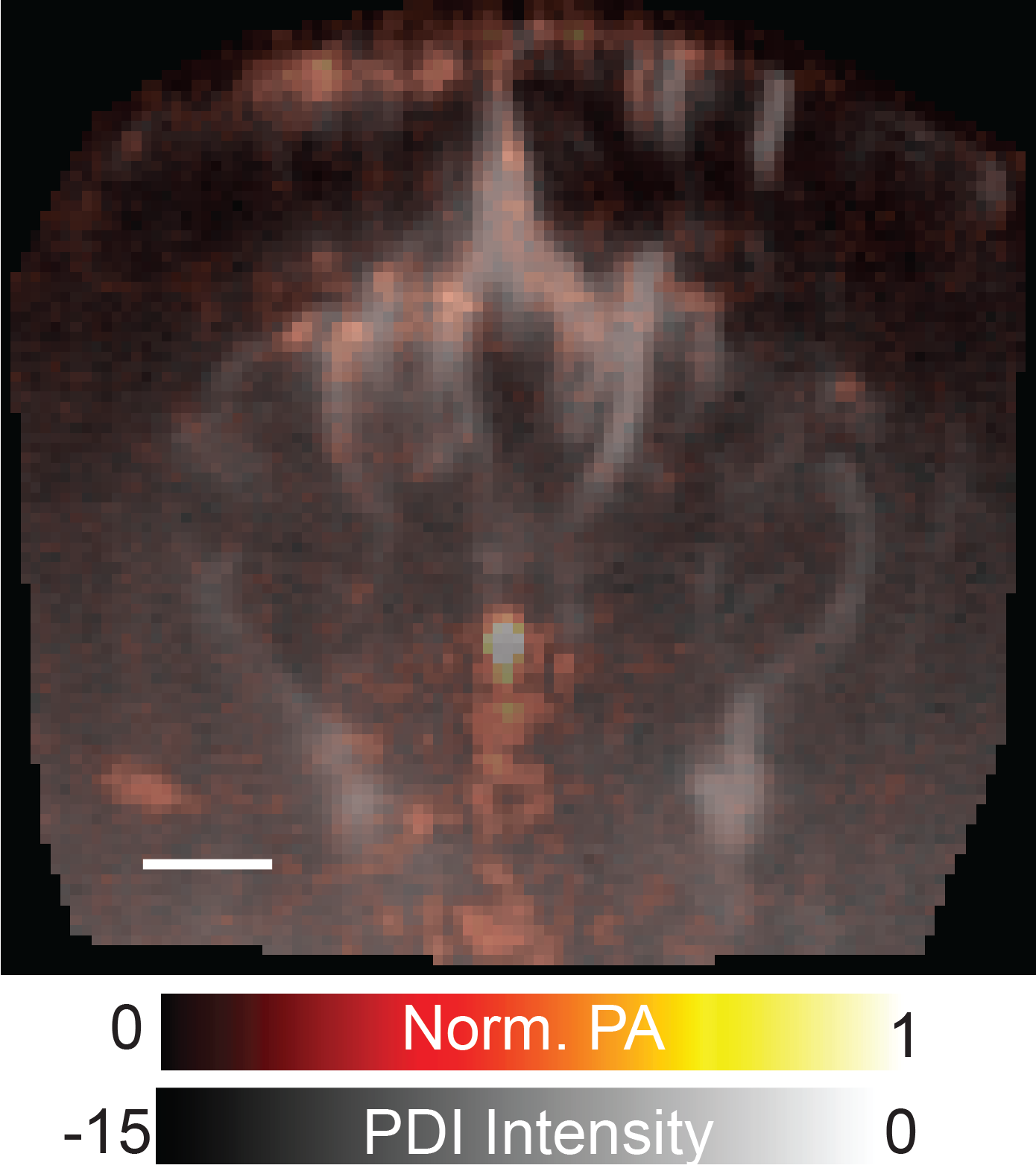

**Supplementary Figure 9: In vivo functional ultrasound and photoacoustic imaging of mouse brain**

**fUS power doppler image in grayscale co-registered with interleaved photoacoustic (PA) B-Mode image in hot colorscale acquired at 800 nm wavelength. While fUS maps cerebral blood volume, PA image contrast represents optical absorption of total hemoglobin at 800 nm.**

**Supplementary Video 1: fUS-optical awake mice imaging**

Video shows fUS-optical awake-mouse acquisition, with the singular-value-decomposition (SVD)-filtered fUS image on the left and the three optical channels on the right. Scale bars, 1 mm.

**Supplementary Video 2: fUS-optical change maps of spontaneous awake mice brain activity**

Video shows a segment of baseline-normalized cerebral blood volume (ΔCBV-fUS), total (ΔHbT), oxy- (ΔHbO), deoxy-hemoglobin (ΔHbR), and calcium (ΔCa²⁺) changes for a mouse in the awake state. Scale bars, 1 mm.

**Supplementary Video 3: fUS-optical change maps of awake mice brain activity during whisker stimulus**

Video shows a segment of baseline-normalized cerebral blood volume (ΔCBV-fUS), total (ΔHbT), oxy- (ΔHbO), and deoxy-hemoglobin (ΔHbR) changes for a mouse in the awake state undergoing whisker stimulation. Scale bars, 1 mm.

**Supplementary Video 4: fUS-optical change maps of spontaneous brain activity in anaesthetized glioblastoma mice at Day 9**

Video shows a segment of baseline-normalized cerebral blood volume (ΔCBV-fUS), total (ΔHbT), oxy- (ΔHbO), deoxy-hemoglobin (ΔHbR), and calcium (ΔCa²⁺) changes for a tumor-bearing mouse at Day 9 post-injection of GL261 glioblastoma cells. Scale bars, 1 mm.

**Supplementary Video 5: fUS-optical change maps of spontaneous brain activity in anaesthetized glioblastoma mice at Day 19**

Video shows a segment of baseline-normalized cerebral blood volume (ΔCBV-fUS), total (ΔHbT), oxy- (ΔHbO), deoxy-hemoglobin (ΔHbR), and calcium (ΔCa²⁺) changes for a tumor-bearing mouse at Day 19 post-injection of GL261 glioblastoma cells. Scale bars, 1 mm.

**Supplementary Video 6: fUS-optical change maps of spontaneous brain activity in anaesthetized glioblastoma mice at Day 23**

### Video shows a segment of baseline-normalized cerebral blood volume (ΔCBV-fUS), total (ΔHbT), oxy- (ΔHbO), deoxy-hemoglobin (ΔHbR), and calcium (ΔCa²⁺) changes for a tumor-bearing mouse at Day 23 post-injection of GL261 glioblastoma cells. Scale bars, 1 mm.
